# *kcnb1* loss-of-function in zebrafish causes neurodevelopmental and epileptic disorders associated with GABA dysregulation

**DOI:** 10.1101/2024.07.03.601913

**Authors:** Lauralee Robichon, Claire Bar, Anca Marian, Lisa Lehmann, Solène Renault, Edor Kabashi, Sorana Ciura, Rima Nabbout

## Abstract

**KEY POINTS:**

- kcnb1 is expressed in distinct cell subtypes and various regions of the central nervous system in zebrafish
- Brain anatomy and neuronal circuits are not disrupted in the *kcnb1* loss-of-function zebrafish model
- Loss of *kcnb1* leads to altered behavior phenotype, light and sound-induced locomotor impairments
- *kcnb1* knock-out zebrafish exhibit increased locomotor sensitivity to PTZ and elevated expression of epileptogenesis-related genes
- *kcnb1^-/-^* larvae show spontaneous and provoked epileptiform-like electrographic activity associated with disrupted GABA regulation

**Objective:** *KCNB1* encodes an α-subunit of the delayed-rectifier voltage-dependent potassium channel K_v_2.1. *De novo* pathogenic variants of *KCNB1* have been linked to developmental and epileptic encephalopathies (DEE), diagnosed in early childhood and sharing limited treatment options. Loss-of-function (LOF) of *KCNB1* with dominant negative effects has been proposed as the pathogenic mechanism in these disorders. Here, we aim to characterize a knock-out (KO) zebrafish line targeting *kcnb1 (kcnb1^+/-^* and *kcnb1^-/-^*) for investigating DEEs.

**Methods:** This study presents the phenotypic analysis of a *kcnb1* knock-out zebrafish model, obtained by CRISPR/Cas9 mutagenesis. Through a combination of immunohistochemistry, behavioral assays, electrophysiological recordings, and neurotransmitter quantifications, we have characterized the expression, function, and impact of this *kcnb1* LOF model at early stages of development.

**Results:** In wild-type larval zebrafish, kcnb1 was found in various regions of the central nervous system and in diverse cell subtypes including neurons, oligodendrocytes and microglial cells. Both *kcnb1*^+/-^ and *kcnb1*^-/-^ zebrafish displayed impaired swimming behavior and “epilepsy-like” features that persisted through embryonic and larval development, with variable severity. When exposed to the chemoconvulsant pentylenetetrazol (PTZ), both mutant models showed elevated locomotor activity. In addition, PTZ-exposed *kcnb1*^-/-^ larvae exhibited higher bdnf mRNA expression and activated c-Fos positive neurons in the telencephalon. This same model presents spontaneous and provoked epileptiform-like electrographic activity associated with disrupted GABA regulation. In this KO model, neuronal circuit organization remained unaffected.

**Significance:** We conclude that *kcnb1* knock-out in zebrafish leads to early-onset phenotypic features reminiscent of DEEs, affecting neuronal functions and primarily inhibitory pathways in developing embryonic and larval brains. This study highlights the relevance of this model for investigating developmental neuronal signaling pathways in *KCNB1*-related DEEs.

## 1. INTRODUCTION

*KCNB1* encodes the pore-forming α1 subunit of the voltagelgated potassium channel subfamily 2 (K_v_2.1). K_v_2.1 is extensively expressed across the central nervous system (CNS), and predominantly localized in clusters on neuronal soma, proximal dendrites, and axonal initial segment of neurons (1–4), generating a delayed-rectifier outward potassium current (1,2,5,6). Moreover, the channel modulates neuronal excitability through activity-dependent regulation of K_v_2.1 phosphorylation in various neuronal subtypes (1,2,5,6). Beyond its role in ionic conduction, K_v_2.1 channels participate in intracellular protein trafficking and calcium signaling (5,7,8).

*De novo* pathogenic variants of *KCNB1* have been reported in patients with developmental and epileptic encephalopathy (DEE), as well as developmental encephalopathy (DE) without epilepsy or with late-onset, severe and pharmacoresponsive epilepsy (9–12). All patients reported in the largest series exhibit poor long-term outcome associated with a wide phenotypic spectrum including severe intellectual disability, attention disorders and autism spectrum disorder (ASD) (9,12,13).

*In vitro* studies of *KCNB1* variants have revealed various degrees of loss-of-function (LOF) exhibiting dominant negative effects such as reduced potassium conductance, altered ion selectivity, and diminished K_v_2.1 channel expression at the cell surface (14–17). These findings are supported by rodent models, including *Kcnb1* knock-out (KO) and knock-in (KI) mice, which exhibit altered behaviors such as locomotor hyperactivity, reduced anxiety-like behavior, and lower seizure thresholds when exposed to chemoconvulsants (18–20). Spontaneous seizures have been reported in homozygous KI *Kcnb1*^R/R^ (*Kcnb1^G379R/G379R^*) (18) and the KI *Kcnb1^R312H^* mouse models (21).

The underlying pathogenic mechanisms of *KCNB1*-related DEEs remain largely uncharacterized in rodent models, and face limitations, particularly regarding drug screening. Zebrafish has gained prominence for studying human neurological disorders, including epilepsy, due to its genetic manipulability, rapid development, and suitability for high-throughput drug screening (22–25). Larval zebrafish, when exposed to the proconvulsant pentylenetetrazol (PTZ), manifest behaviors akin to seizures, making them a valuable model for epilepsy investigations (26) such as Dravet syndrome related to *SCN1A* (26–28) and epilepsies associated with *DEPDC5* (29,30). In addition, drug screening in larval zebrafish has been performed in both genetic and chemically induced epilepsy models, demonstrating the translational potential of this model by identifying compounds capable of reducing the severity of epileptic seizures (31–34).

The zebrafish orthologue of *KCNB1, kcnb1,* shares 67% sequence homology with the human gene. The expression of *kcnb1* starts at 19 hours post-fertilization (hpf), increases during zebrafish development, and is mainly localized in the brain of adult zebrafish with tissue-specific expression patterns comparable to mammals (35). The first *kcnb1* KO zebrafish model (*kcnb1*^-/-^) was generated using CRISPR/Cas9 mutagenesis, leading to a premature stop codon between the N-terminal cytoplasmic domain and the first transmembrane domain of the protein (36). This *kcnb1*^-/-^ model revealed developmental abnormalities such as a small proportion of gastrulation defect and reduced brain ventricles (36) along with disruption of inner ear development (35), emphasizing *kcnb1*’s role in developmental signaling pathways.

However, the effects of *kcnb1* loss-of-function on brain in development, in the context of epilepsy, remain unexplored. In this study, we aim to characterize the behavioral, electrophysiological and molecular consequences of this *kcnb1*^-/-^ zebrafish model at various developmental stages. This analysis will shed light on the key features of *KCNB1*-related DEEs, contributing to a better understanding of the pathophysiological mechanisms underlying this disorder.

## 2. MATERIALS AND METHODS

### 2.1. Housing conditions of zebrafish

All procedures were approved by the Institutional Ethics Committees at the Research Centers of IMAGINE Institute (INSERM U1163, Paris, France) and were performed in accordance with the European Union Directive (2010/63/EU). Adult zebrafish (Danio Rerio) were housed in a conventional animal facility. Experiments were performed on wild-type AB and TU strains (Tübingen, Germany) and *kcnb1* mutant zebrafish lines (*kcnb1*^+/-^ and *kcnb1*^-/-^) staged from 0-to 6-days post-fertilization (dpf) (37). Fertilized eggs were collected by natural spawning. Embryos and larvae were maintained at 28 ± 1 °C in a non-CO_2_ incubator (VWR) with a 14h/10h light/dark cycle in embryo medium (Instant Ocean).

### 2.2. kcnb1 mutant zebrafish lines

Heterozygous *kcnb1* mutant embryos (*kcnb1*^+/-^) were provided by Dr. Vladimir Korzh (Warsaw, Poland). The *kcnb1* mutant line was generated using CRISPR/Cas9 mutagenesis, resulting in a premature stop codon (36). *kcnb1*^+/-^ mutants were raised to adulthood and crossed to produce *kcnb1*^-/-^ zebrafish for experiments (*kcnb1^sq301/sq301^*, ZDB-ALT-170417-2) (36). Genomic DNA was extracted from adult zebrafish fins. Samples were amplified by PCR (Bio-Rad) using DreamTaq Hot Start PCR Master Mix (Thermo Scientific) and primers targeting *kcnb1* (10 mM): F_5’-TGTGACGACTACAACCTGGA-3’ and R_5’-CTCCTCGTTCATCTGCTCCT-3’. DNA samples were sent for sequencing to GATC Biotech, Eurofins Genomics.

### 2.3. Survival assay and morphological analysis

Zebrafish embryos and larvae were maintained at 28 ± 1 °C non-CO_2_ incubator (VWR), and were fed daily from 0 to 15 dpf. After 6 dpf, larvae were transferred to tanks with dripped water flux. Survival was assessed based on the number of living individuals relative to the total population. Zebrafish were photographed using a stereomicroscope (SZX16, Olympus Life Science). Measurements of the body length and the head surface were performed manually with ImageJ software (National Institutes of Health, NIH).

### 2.4. Reverse transcription quantitative polymerase chain reaction (RT-qPCR)

cDNA from larval zebrafish at 6 dpf (30 per pool) was synthesized using 5X All-In-One RT MasterMix (abm, Canada). qPCR on *kcnb1* and *bdnf* was performed with BlasTaq 2X qPCR Master Mix (abm) on a BioRad CFX384 System (Bio-Rad). Relative gene expression was determined by the 2^-ΔΔCt^ method, normalized to β*-actin* or *ef1*α, with *kcnb1*^+/+^ serving as the reference (relative fold change = 1). The following primers were used for qPCR: β*-actin* (F_5’-CGAGCTGTCTTCCCATCCA-3’, R_5’-TCACCAACGTAGCTGTCTTTCTG-3’); *ef1*α (F_5’-CTGGAGGCCAGCTCAAACAT-3’, R_5’-ATCAAGAAGAGTAGTACCGCTAGCATTAC-3’); *kcnb1 (F_*5’-TGAAGTTCCGGGAGAGTGTT-3’, R_5’-CAGGTTGGCGATGTCGTTCT-3’); *bdnf (F_*5’-GACTCGAAGGACGTTGACCTGTA-3’, R_5’-CGGCTCCAAAGGCACTTG-3’).

### 2.5. Immunoprecipitation (IP) and western blot of kcnb1

Tissues from larval zebrafish at 6 dpf (30 per pool) were homogenized in lysis buffer supplemented with protease inhibitors (Roche). Lysates were centrifugated at 12,000 × g for 15 minutes at 4°C followed by a BCA Protein Assay Kit (Invitrogen). Immunoprecipitation was performed to detect kcnb1 by using the Immunoprecipitation Kit dynabeads protein G (Invitrogen), following the manufacturer’s protocol. 200 µg of the supernatant with 4 µg of anti-kcnb1 (ProteinTech, #19963-1-AP) or anti-Pan actin (Thermo Scientific, #ms-1295) were used and IgG antibody served as a negative control. Membranes from western blotting were detected using an ECL reagent (Cytiva) and the ChemiDoc™ Imaging Systems (Bio-Rad). Densities of kcnb1 bands were normalized to Pan-actin expression using ImageJ (NIH).

### 2.6. Locomotion assessment

#### Premotor activity

Tail coiling activity in zebrafish embryos was recorded every hour from 24 hpf to 36 hpf. Sixty-second movies were obtained under a stereomicroscope with darkfield illumination (SZX16, Olympus Life Science), recorded at 30 frames per second (fps). Premotor activity was counted manually using ImageJ software (NIH).

#### Touch-evoked escape response (TEER)

The tail of 48 hpf-embryos was mechanically stimulated, and their swimming trajectories were recorded at 30 fps using SpinView V2.0.0.147 software (FLIR Systems lnc.). The trajectory was traced using the Manual Tracking Plug-in in ImageJ software (NIH), and analyzed for distance, velocity, and time spent in motion (see **Figure 3A**).

#### Spontaneous locomotion and pentylenetetrazol (PTZ)-induced seizures

Larvae were acclimatized in a 48-well plate (Fisher Scientific) with lights off for 15 min into a Zebrabox equipped with Zebralab 438 software (Viewpoint Life Sciences). Larvae were submitted to two different protocols: 1) a light-induced protocol of 60 minutes recording split into 10 min light/dark conditions repeated three times and 2) a sound-induced protocol of four repeated audio stimuli (450 hertz, 80 decibels, 1 second/audio). The 5 seconds following audio stimuli were analyzed. To evaluate the effect of PTZ on locomotor activity, larvae were first recorded for 30 min to determine baseline activity levels. Fresh 5 mM PTZ (Sigma-Aldrich) was then added and zebrafish were recorded for a further 30 min (see **Figure 4A**).

### 2.7. Electrophysiological analysis

Larval zebrafish at 5 and 6 dpf were embedded in 1% low-melting-point agarose (Sigma-Aldrich) and covered with artificial cerebrolspinal fluid (ACSF, pH 7.8, osmolarity to 290-295 mOsm/l). A microelectrode (2–7 MΩ) was filled with ACSF and implanted into the optic tectum, using an Olympus microscope (Infinity 3S, 10X magnification). Local field potential (LFP) recordings were obtained using a MultiClamp700B amplifier (Molecular devices) coupled with an Axon Digidata 1550 (Molecular devices) and the Clampex V11.1 software (Molecular devices). Baseline recordings of 30 min duration were assessed to larvae before adding 40 mM PTZ (Sigma-Aldrich). After 5 min of treatment, the recording continued for 30 min. Negative spikes were automatically analyzed using Clampfit V11.2 software (Molecular devices) during a 10 min interval, using a low-pass Gaussian filter of 560 Hz and a digital reduction of 10 (see **Figure 5A**).

### 2.8. Immunohistochemistry on slices

Zebrafish aged from 48 hpf to 6 dpf were fixed as previously described (30). Brain slices of zebrafish (20 µm) were obtained using the cryostat CM3050S (Leica). After permeabilization for 30 minutes, the slices were incubated overnight at 4°C in blocking solution containing primary antibodies: Kcnb1 (1:100, Tebubio, #PAB7569), NeuN (1:100, Merck, #ABN90), Olig2 (1:100, DSHB, #PCRP-OLIG2-1E9-s), CX3CR1 (1:100, ProteinTech, #13885-1-AP), Ankyrin G (1:100, Proteintech, #27980-1-AP) and HuC (1:100, Tebubio, #FNab04072). Secondary antibodies (Alexa Fluor 488, Alexa Fluor 568 and Alexa Fluor 647, Thermo Scientific) were incubated in blocking solution for 2h at room temperature (RT). Slices were incubated in DAPI (Invitrogen, #D3571) followed by mounting on glass slide in Immu-Mount mounting medium (Epredia, #9990402). Images were captured using a Spinning disk Zeiss system (Carl Zeiss). Colocalization, represented in white, was determined using Z-stack projection on IMARIS V10.1.0 software (Oxford Instruments).

### 2.9. Whole-mount immunohistochemistry

Whole-mount immunohistochemistry on 48 hpf and 6 dpf zebrafish was performed according to a previously established protocol (38). Embryos were treated with 0,003% 1-Phenyl-2-thiourea (PTU) (Sigma-Aldrich) to prevent pigmentation of the skin. Briefly, zebrafish were fixed with 4% formaldehyde (Sigma-Aldrich) for 2h at RT. A dehydration with methanol (Sigma-Aldrich) was followed by a gradual rehydration. Zebrafish were permeabilized in fresh acetone (Sigma-Aldrich) for 20 min and blocked with 10% Bovine standard-albumin (BSA) (Eurobio Scientific) overnight at 4°C. They were incubated with the primary antibody (1% BSA / 0,1%PBST) at 4°C for two days. Primary antibodies used at 1:100 were: acetylated tubulin (Sigma-Aldrich, #T7451), 3A10 (DSHB, #3A10) and c-Fos (Santa Cruz, #sc-166940). Secondary antibodies were incubated in 1% BSA / 0,1% PBST overnight at 4°C using Alexa Fluor 488 and Alexa Fluor 647 (Thermo Scientific) at 1:250. After passing through increased percentages of glycerol (Sigma-Aldrich), image acquisition was performed under a Spinning disk Zeiss system (Carl Zeiss). The same parameters were applied to all images using ImageJ (NIH).

### 2.10. GABA and Glutamate ELISA assays

Based on the PTZ treatment protocol, the head of 6 dpf larvae (50 per pool) was collected at the end of each 30 min-period recording of the basal activity and 5 mM PTZ treatment (see **Figure 5A**). Same reagents were used as the IP protocol for tissue homogenizing. Enzyme-linked immunosorbent assay (ELISA) was performed to quantify gamma-aminobutyric acid (GABA, immusmol, #BA E-2500) and glutamate (immusmol, #BA E-2400) from 20 µg of supernatant, following the manufacturer’s protocol.

### 2.11. Statistical analysis

Statistical analysis was performed using GraphPad Prism V.8 software (GraphPad Software). Statistical tests used are specified in the legend of each figure. Values represent the mean ± standard error of the mean (SEM). Results were significant when the p-value was < 0.05. For all experiments, at least 3 single experiments were performed (N) with a certain number of embryos or larvae per condition (n).

## 3. RESULTS

3.1. *kcnb1 is expressed in diverse cell-subtypes and regions of the CNS in Wild-type larval zebrafish*

Previous studies have shown significant expression of *kcnb1* in the eyes, ears, and central nervous system (CNS) of wild-type (WT) larval zebrafish using whole-mount *in situ* hybridization (36). To further investigate the distribution of the kcnb1 protein in the brain, we performed immunohistochemistry on WT larvae at 6 dpf **(Figure 1A-1F, Supporting Information 1A-1B)**. Our findings revealed broad expression of kcnb1 throughout multiple regions of the CNS, such as the diencephalon, midbrain, telencephalon and hindbrain **(Figure 1A-1C, Supporting Information 1A)** including the spinal cord **(Figure 1C)**, although the protein was difficult to detect in the eyes. Further analysis demonstrated that kcnb1 is expressed in various CNS cell subtypes. Notably, kcnb1 is localized in neurons as evidenced by colocalization with NeuN, a neuronal nuclear marker, and further confirmed by Ankyrin G marker, expressed on the axonal initial segment of neurons **(Figure 1D, Supporting Information 1B)**. kcnb1 was also found in oligodendrocytes, marked by Olig2, a transcription factor **(Figure 1E)**, and in microglial cells, identified by CX3CR1, a fractalkine receptor marker **(Figure 1F)**. These results provide the first characterization of kcnb1 expression in distinct CNS cell subtypes in zebrafish.

**Figure 1.**
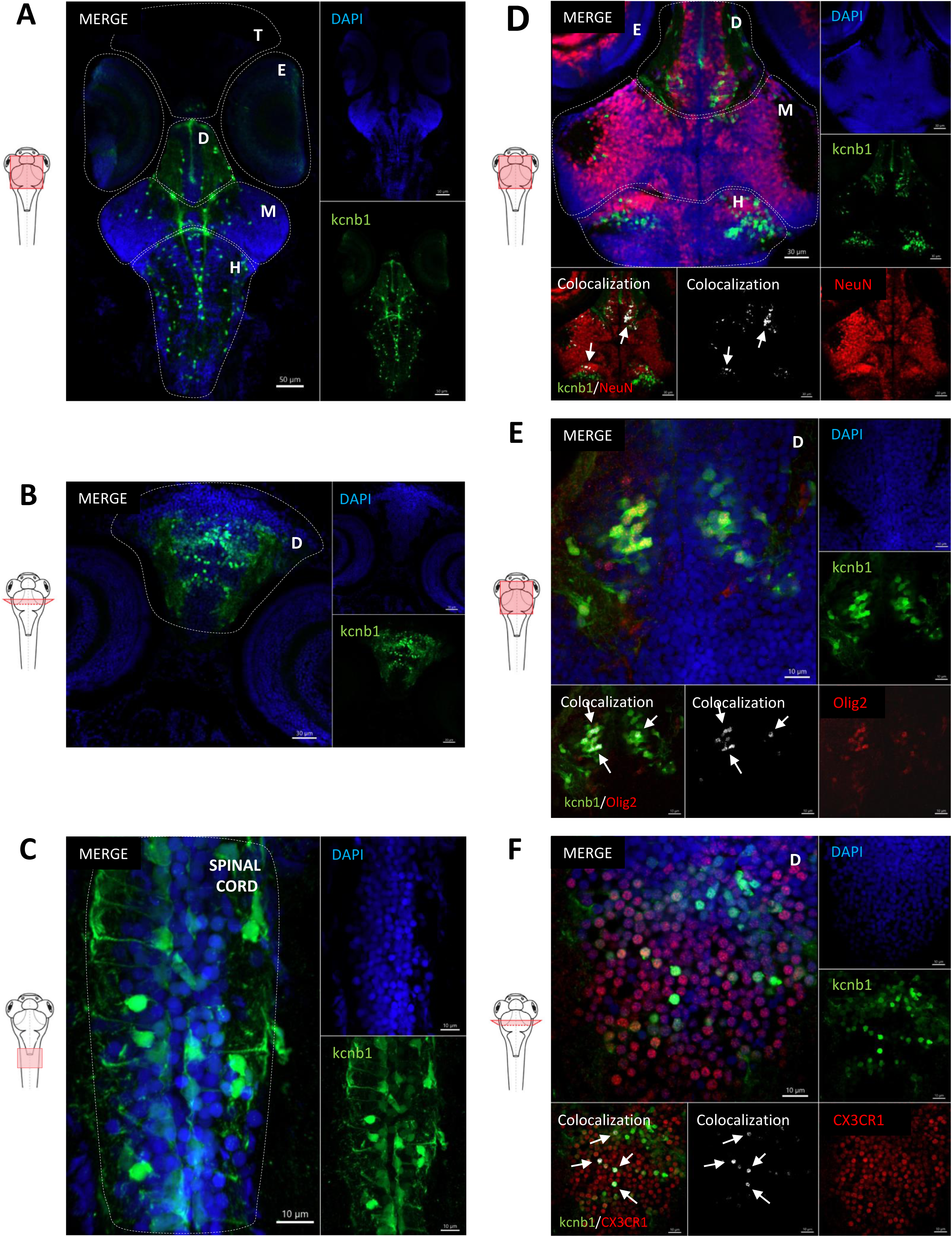
kcnb1 is expressed in distinct cell subtypes and various regions of the central nervous system in 6 dpf Wild-type zebrafish. A and B. Horizontal and transversal sections of Wild-type (WT) zebrafish expressing cells (DAPI, blue) labelled with anti-kcnb1 (green), showing a large expression of the protein in the central nervous system (CNS) at 6-day post-fertilization (dpf). The protein is expressed in the telencephalon, diencephalon, midbrain including the optic tectum, and the hindbrain comprising the cerebellum and the spinal cord (**A:** scale bar 50 µm; magnification10x; **B:** scale bar 30 µm; magnification 20x). **C.** Horizontal section of a WT zebrafish at 6 dpf showing the presence of kcnb1 along the spinal cord (scale bar 10 µm; magnification 63x). **D-F.** Horizontal sections of WT zebrafish expressing cells (DAPI, blue), kcnb1 (green) and specific cell subtype markers, respectively **(D)** a neuronal nuclear marker (NeuN, red, scale bar 30 µm; magnification 20x); **(E)** an oligodendrocyte transcription factor 2 (Olig2, red, scale bar 10 µm; magnification 63x) and **(F)** CX3C motif chemokine receptor 1 expressed in microglial cells (CX3CR1, red, scale bar 10 µm; magnification 63x). Images demonstrate the colocalization between kcnb1 and the 3 different cell subtypes markers, represented in white and marked by arrows. These results indicate the presence of kcnb1 in neurons, oligodendrocytes and microglial cells. Colocalization was determined using Z-stack projection on IMARIS V10.1.0 software (Oxford Instruments). In figures, CNS regions are delimited by dotted lines. CX3CR1: CX3C motif chemokine receptor 1; D: Diencephalon; DAPI: 4,6ldiamidinol2lphenylindole; E: Eyes; H: Hindbrain; kcnb1: Potassium voltage-gated channel subfamily B member 1; M: Midbrain; NeuN: Neuronal nuclear antigen; Olig2: Oligodendrocyte transcription factor; T: Telencephalon. n = 3-4 fish/section.

### 3.2. kcnb1 knock-out expression and morphological analyses

Using a *kcnb1* knock-out zebrafish model generated by Shen et collaborators (2016), we investigated loss-of-function effects of *kcnb1* in the context of DEEs. We first confirmed the genomic sequence of *kcnb1*^-/-^ fish by Sanger sequencing, identifying a distinct 2-G bases deletion along with a 14-bp insertion on chromatograms (Δ14bp) **(Figure 2A)**. Realltime qPCR analysis revealed a significant reduction in *kcnb1* transcript expression by 44% in *kcnb1*^+/-^ and by 56% in *kcnb1*^-/-^ zebrafish larvae compared to WT at 6 dpf **(Figure 2B)**. Despite this reduction in transcript levels, kcnb1 protein expression remained comparable across all genotypes **(Figure 2C)**. The survival rates of both *kcnb1^+/-^ and kcnb1^-/-^* lines from 0 to 15 dpf were similar to those of WT **(Figure 2D and Table 1)**. Measurements of body length and head surface of *kcnb1* mutants (*kcnb1*^+/-^ and *kcnb1*^-/-^) showed no significant morphological differences compared to WT at 48 hpf and 6 dpf **(Figure 2E-2G)**, corroborated by using the pan-neuronal marker HuC **(Supporting information 1C)**. Visualization of neuronal fibers with an acetylated tubulin marker, revealed no major neuronal loss or organizational defects due to *kcnb1* loss-of-function during early development (**Figure 2H**), supported by similar Mauthner cell body length observed across conditions at 48 hpf (M-cell, 3A10 marker, **Supporting information 1D-1E**). These results indicate that *kcnb1* loss-of-function does not impact the normal growth of fish during early development.

**Figure 2.**
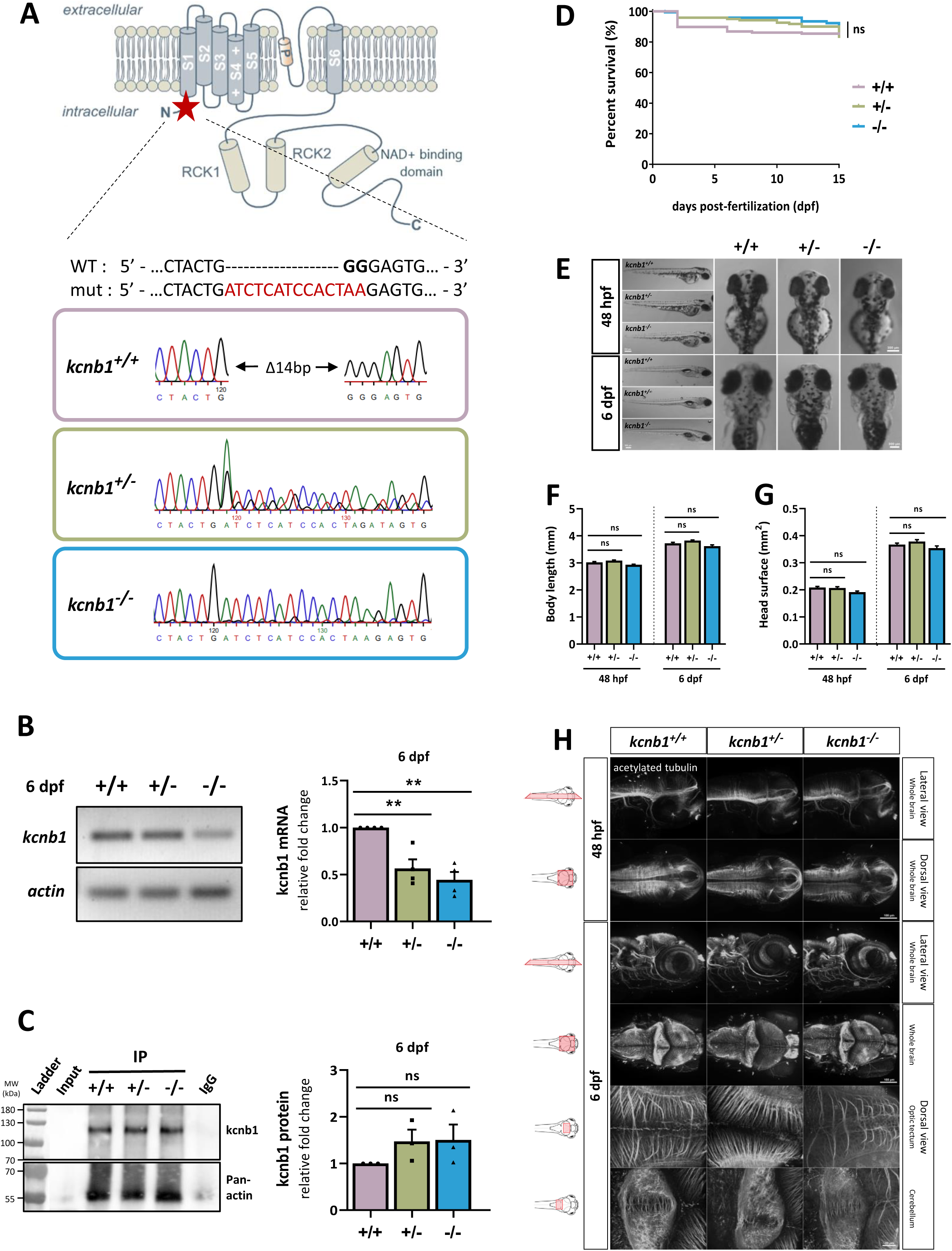
Genotypic characterization of a *kcnb1* knock-out zebrafish model that is not affected by brain anatomical defects. **A.** Schematic illustration of the generation of a *kcnb1* knock-out zebrafish model obtained by Shen et al. (2016). The representation of the α-subunit *kcnb1* is adapted from a previous schematic picture (41). Insertion of a 14 base-pairs (bp)-nucleotide sequence (identified in red) and of a 2 bp-nucleotide deleted (identified in black bold type) into the first exon of the gene introducing a premature stop (Δ14bp) between the N-terminal region and the first transmembrane domain of the protein, using the CRISPR-Cas9 system (targeted sequence: GGAGCTGGACTACTGGGGAG in *kcnb1* exon 1; ID zfin: ZDB-ALT-170417-2; indel mutation; line: Kcnb1^sq301/sq301^). **B.** RT-qPCR analysis of total *kcnb1* at 6 dpf demonstrating a significant decrease of *kcnb1* mRNA in *kcnb1^+/-^* and *kcnb1^-/-^*compared with *kcnb1^+/+^* fish (N = 4; n = 30/sample; One-Way ANOVA with Bonferroni post-hoc test; **p<0,01). Data are normalized to actin mRNA expression and the condition *kcnb1^+/+^* is considered as the reference value (relative fold change = 1). **C.** Immunoprecipitation (IP) of kcnb1 in 6 dpf zebrafish demonstrating similar profile of protein expression between the three genotypes (N = 3; n = 30/sample; One-Way ANOVA with Bonferroni post-hoc test; ns: non signifcant). Input was used as a control of Western blot and IgG antibody served as a negative control of IP. Data are normalized to Pan-actin protein expression, with *kcnb1*^+/+^ serving as the reference (relative fold change = 1). **D.** Kaplan-Meier survival curve between 0 and 15 dpf showing that the partial or complete loss of *kcnb1* do not impact the normal growth of fish at early stages of development (see **Table 1**; N = 3 repeats; n = 121-169/genotype; Log-rank test; ns: non significant). **E.** Images showing that *kcnb1^+/-^*and *kcnb1^-/-^* embryos and larvae do not show gross morphological changes at 48 hours post-fertilization (hpf) and 6 dpf (scale bar: 300 µm). **F-G.** Quantification of major morphological aspects at 48 hpf and 6 dpf including measurement of **(F)** body length and **(G)** head surface. The *kcnb1* mutant zebrafish models (*kcnb1^+/-^* and *kcnb1^-/-^*) do not show any significant change in each parameter at different developmental stages as compared to *kcnb1^+/+^* (N = 4 repeats; n = 28-38 ZF/genotype; one-Way ANOVA with Bonferroni post-hoc test; ns: non-significant). **H.** Whole-mount images of embryonic (48 hpf) and larvae (6 dpf) zebrafish immunostained with anti-acetylated tubulin marker to identify global circuits of neuronal fibers (lateral and dorsal view; 3D reconstruction; magnification 20x and scale bar at 100 μm; magnification 40x and scale bar at 20 µm). The partial or complete loss of *kcnb1* does not affect the neuronal brain density at different early stages of development (n = 5-7 ZF/genotype/developmental stage).

**Table 1:**
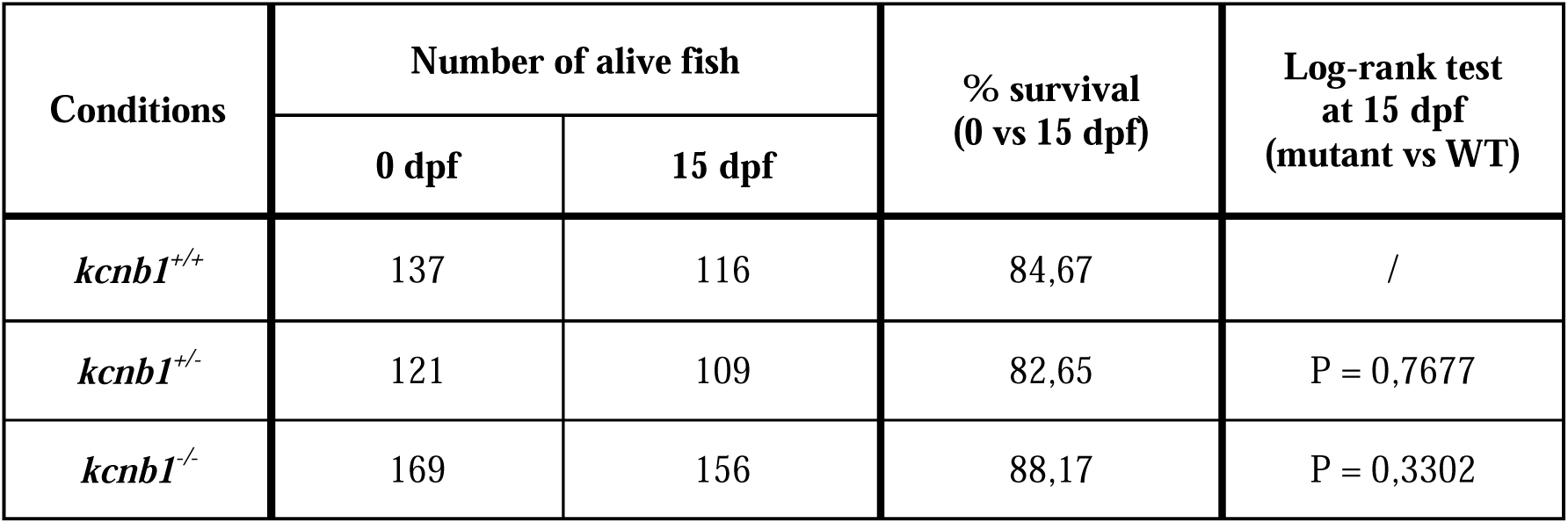
Survival table between 0 and 15 days post-fertilization (dpf). Results show a similar growth of *kcnb1* mutant fish (*kcnb1^+/-^*and *kcnb1^-/-^*) as compared to control fish (*kcnb1^+/+^*) at early stages development (N = 3 repeats; n = 121 - 169 ZF/genotype, Log-rank test, ns: non-significant).

### 3.3. Loss of kcnb1 leads to altered behavior phenotype, light and sound-induced locomotor impairments

*KCNB1* mutations in patients are associated with a range of locomotor disabilities including hyperactivity, myoclonia, ataxia and hypotonia (9). First evidence of motor impairments was observed in both *kcnb1* mutants showing a drastic decreased of tail coiling activity compared to WT zebrafish from 24 to 36 hpf **(Supporting information 2A-2B)**. Using the touch-evoked escape responses (TEER) test at 48 hpf **(Figure 3A)**, *kcnb1*^+/-^ and *kcnb1^-/-^* zebrafish exhibited rapid circular swimming, in contrast to the straight-line swimming observed in WT condition **(Figure 3B)**, significantly increased in terms of total distance swam, velocity and time spent in motion **(Figure 3C-3E)**. Notably, within the same condition, we observed a variability in the swimming behavior of embryos classified into two groups: those with a severe phenotype (completing a minimum of two swim circles) and those with a mild phenotype (grouping other trajectories) **(Table 2**). We found that 30% of *kcnb1*^+/-^ embryos displayed a severe phenotype compared to 52% of *kcnb1*^-/-^ zebrafish, with an increase of studied parameters **(Table 2, Supporting information 2C-2F)**. While all three conditions showed low locomotor activity in response to light changes at 3 and 4 dpf, this activity significantly increased in all conditions at later stages **(Figure 3H)**. However, starting at 5 dpf, *kcnb1^-/-^* larvae displayed pronounced locomotor hyperactivity compared to WT and *kcnb1^+/-^* zebrafish **(Figure 3H)**. This finding is aligned with results from **Figure 3G and 3F**, showing significant locomotor hyperactivity in response to light/dark transitions. In a second protocol, both *kcnb1^+/-^*and *kcnb1*^-/-^ mutants exhibited a significant decrease in locomotor activity after each audio stimulus compared to the WT condition **(Figure 3I)**. These results suggest that the locomotor phenotype may be due to dysregulation of electrical neuronal activity affecting different sensorimotor pathways.

**Figure 3.**
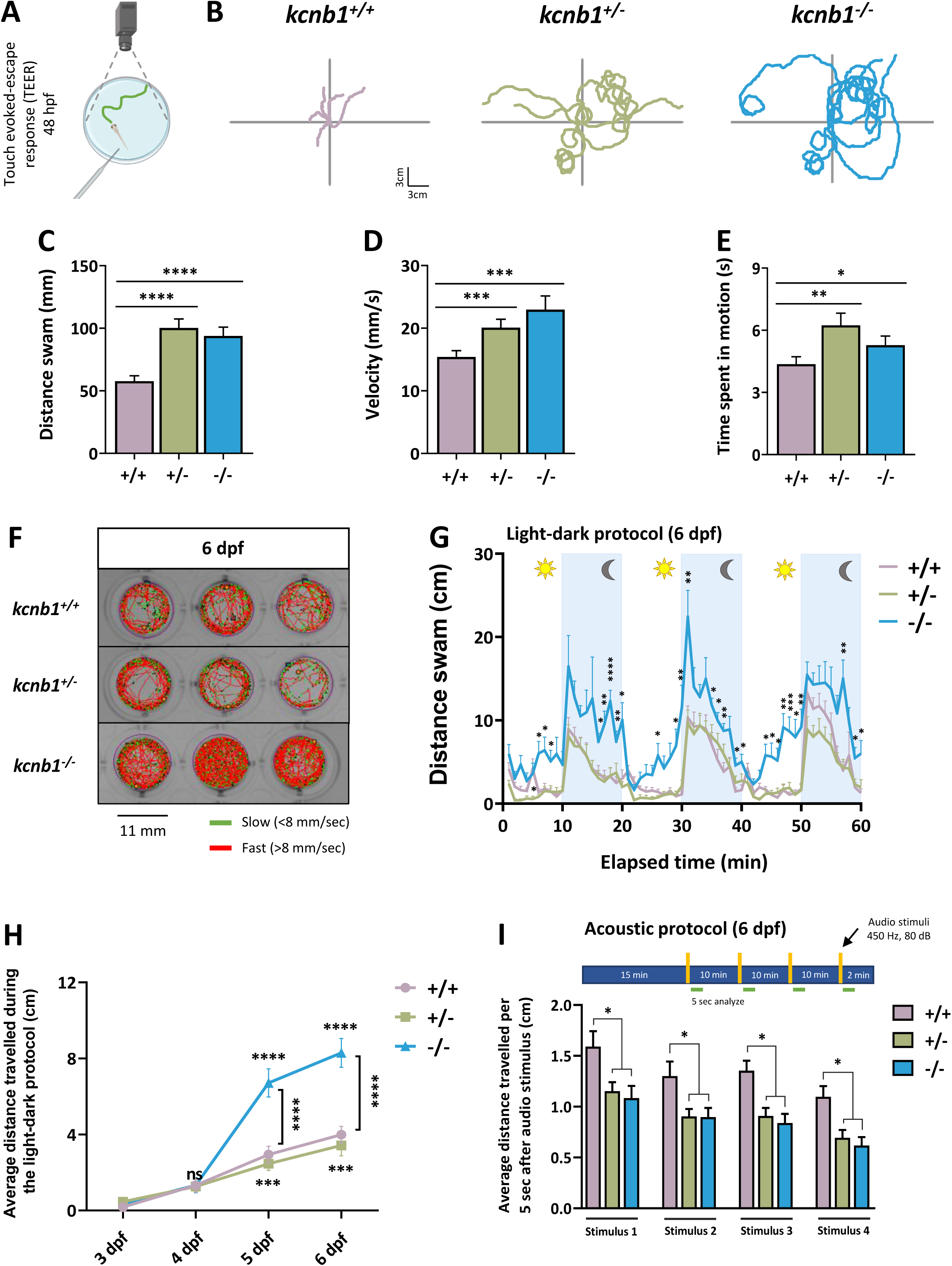
Loss of *kcnb1* leads to altered behavior phenotype, light and sound-induced locomotor impairments. **A.** Schematic representation of the touch evoked-escape response (TEER) test. The tail of 48 hpf-embryos is mechanically stimulated, and the swimming trajectory of zebrafish is recorded. **B.** Representative traces of individual swimming episodes at 48 hpf showing the typical tortuous trajectory of *kcnb1* mutant models (*kcnb1^+/-^* and *kcnb1^-/-^*) as compared to the straight-line trajectory of *kcnb1^+/+^* zebrafish (5 trajectories/genotype). **C-E**. The quantification of the swimming trajectory tortuosity shows a significant increase of different studied parameters in both *kcnb1* mutant conditions as compared to WT zebrafish, including an increase of **(C)** the distance swam, **(D)** the velocity and **(E)** the time spent in motion. Furthermore, within the same genotype, a huge variability in the swimming behavior has been observed and fish were divided into two distinct phenotypes: severe (at least two swim circles) and mild (other trajectories). The subdivision of both phenotypes is represented in **Table 2** and **Supporting information 2C-2F** (N = 3 repeats; n = 50-88 ZF/genotype; one-way ANOVA with Bonferroni post-hoc test; ns: non-significant; *p<0,05; **p<0,01; ***p<0,001; ****p<0,0001). **F.** Schematic representation of individual trajectory of 3 larvae zebrafish per genotype obtained with ViewPoint software (Zebrabox). Between 3 and 6 dpf, a protocol was applied to fish with 3 repetitions of 10 minutes in the light followed by 10 minutes in the dark (green lines: slow movements <8 mm/sec; red lines: fast movements >8 mm/sec). **G.** Average distance swam by larvae zebrafish at 6 dpf during the whole light-dark protocol described previously. Spontaneous locomotor hyperactivity was observed in *kcnb1^-/-^*larvae in the light and maintained significantly increased during the dark phase as compared to WT and *kcnb1^+/-^* larvae (N = 3 repeats; n = 35 ZF/genotype; one-way ANOVA with Bonferroni post-hoc test; *p<0,05; **p<0,01; ***p<0,001; ****p<0,0001). **H.** Representation of average distance travelled by larvae at different days of development (from 3 to 6 dpf) following the 10 min light-dark protocol descrived. At 3 and 4 dpf, controls and *kcnb1* mutant fish (*kcnb1^+/-^* and *kcnb1^-/-^*) present low locomotor activity that was significantly increased from 5 to 6 dpf. A significant gap was observed starting from 5 dpf with a locomotor hyperactivity for *kcnb1^-/-^* larvae as compared to WT and *kcnb1^+/-^* fish, reflecting the result obtained in Figure 3G (N = 3 repeats; n = 35 ZF/genotype; one-way ANOVA with Bonferroni post-hoc test; the locomotor activity of zebrafish from the same genotype was compared from 4 to 6 dpf with a reference value at 3 dpf and the three genotypes were also compared between them; ns: non-significant; ***p<0,001; ****p<0,0001). **I.** Quantification of the distance swam during the 5 seconds following each of the four audio stimuli applied (450 Hertz; 80 decibel; 1 second) to larvae at 6 dpf. Both mutant conditions (*kcnb1^+/-^* and *kcnb1^-/-^*) present a significant decrease of the locomotor activity in response to audio stimuli as compared to the WT condition (N = 3 repeats; n = 48-64 ZF/genotype; one-way ANOVA with Bonferroni post-hoc test; *p<0,05).

**Table 2:**
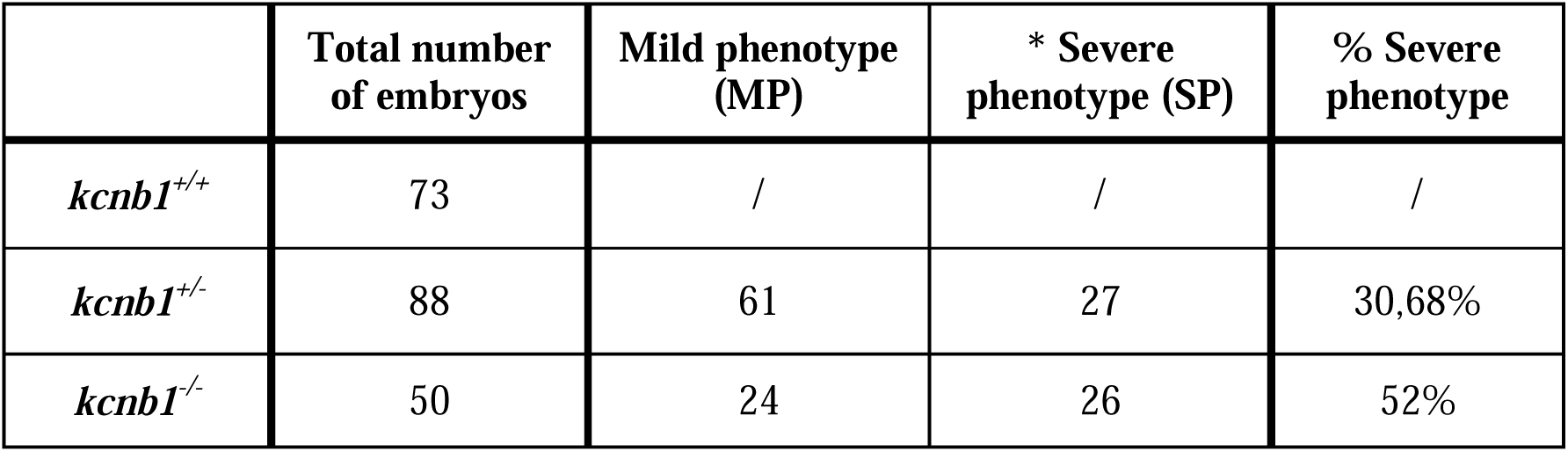
Diverse phenotypes were observed during the Touch-Evoked Escape Response (TEER) of 48 hpf-mutant embryos. Within the same genotype, embryos were divided in severe phenotype (*: swimming trajectory with at least two swim circles; SP) and mild phenotype (other trajectories; MP) indicating a huge variability in swimming behavior after a TEER test of *kcnb1^+/-^* and *kcnb1^-/-^* fish (N = 3 repeats; n = 50-88 ZF/genotype).

### 3.4. kcnb1^-/-^ zebrafish exhibit increased sensitivity to PTZ-induced seizures and elevated expression of epileptogenesis-related genes

Previous studies have shown that exposure of zebrafish larvae to pentylenetetrazol (PTZ) increases seizure-like behavior in a concentration-dependent manner (26). We recorded the baseline locomotor activity of 6 dpf larvae for 30 minutes, followed by a 30-minute exposure to 5 mM PTZ **(Figure 4A)**. Low baseline locomotor activity of *kcnb1* mutants was similar to that of WT larvae **(Figure 4B-4C)**. However, after PTZ exposure, *kcnb1* mutants (*kcnb1^+/-^* and *kcnb1*^-/-^) exhibited significantly higher swimming activity, characterized by fast circles **(Figure 4B-4C)**. To explore potential molecular disruptions, we measured *bdnf* (Brain-Derived Neurotrophic Factor) mRNA expression, a neurogenesis and epileptogenesis-related gene, before and after PTZ treatment **(Figure 4A and 4D)**. *bdnf* expression was similar between conditions during baseline activity, although significantly increased after PTZ treatment in the *kcnb1^-/^*^-^ condition as compared to *kcnb1^+/-^* and WT zebrafish **(Figure 4D)**. In addition, we assessed c-Fos expression, an early gene marker of epileptic seizures, in the telencephalon of 6 dpf larvae **(Figure 4A, 4E-4G, Supporting information 3)**. The number of c-Fos-positive neurons was similar across all conditions, both before and after PTZ treatment **(Figure 4F)**. We analyzed the distribution of activated neurons according to the fluorescence intensity value of c-Fos, reflecting the level of neuronal activation divided in: low (0-25%), moderately low (25-50%), moderately high (50-75%) and high (75-100%) neuronal activation **(Figure 4G, Supporting information 3B)**. *kcnb1^-/^*^-^ fish showed a shift towards higher levels of neuronal activation, with a significant increase in moderately low and moderately high activation levels after PTZ treatment **(Figure 4G, Supporitng information 3B)**. These results suggest that the *kcnb1* LOF model exhibits neurogenesis impairments in the developing brain of zebrafish.

**Figure 4.**
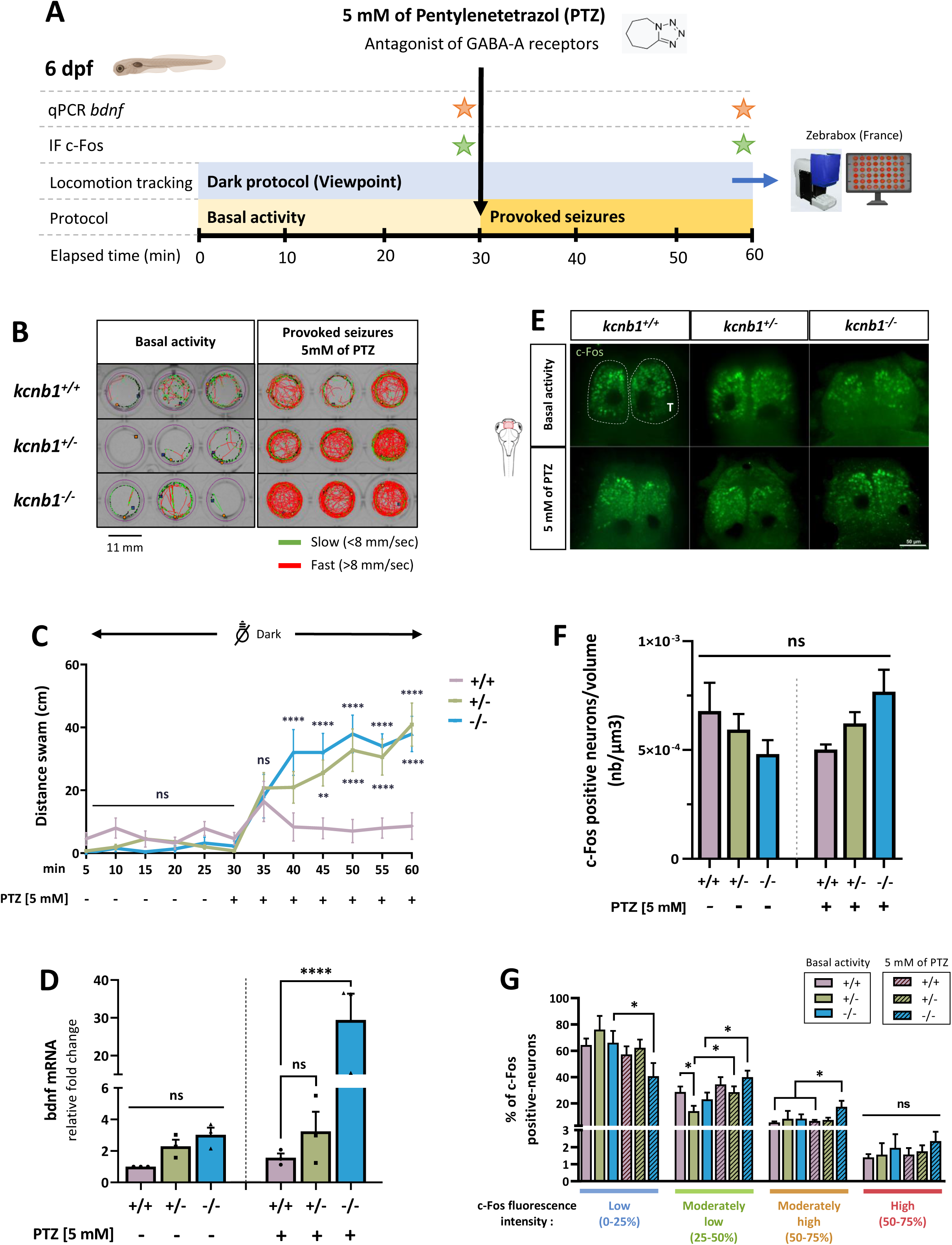
*kcnb1* knock-out zebrafish exhibit increased locomotor sensitivity to PTZ and elevated expression of epileptogenesis-related genes. A. Schematic representation of the protocol followed to identify impact of PTZ on locomotion, bdnf and c-Fos expression. To test seizure susceptibility in the *kcnb1* LOF model, we recorded the baseline locomotor activity of 6 dpf larvae for 30 minutes, followed by a 30-minute exposure to 5 mM PTZ (Zebrabox, ViewPoint). We then quantified two epileptogenesis-related genes: *bdnf* (Brain-Derived Neurotrophic Factor) by qPCR and c-Fos by immunofluorescence. **B.** Schematic representation of individual trajectory of 3 larvae zebrafish per genotype obtained with ViewPoint software (Zebrabox). Fast swimming circles were observed for both mutant larvae conditions (*kcnb1^+/-^* and *kcnb1^-/-^*) after 30 minutes of 5 mM Pentylenetetrazol (PTZ) treatment, a pro-convulsant. Green lines: slow movements (<8 mm/sec), red lines: fast movements (>8 mm/sec). **C.** Global locomotor activity of 6 dpf-zebrafish into the dark was recorded by applying a protocol of 30 minutes of basal activity followed by 30 minutes of chemically induced-seizures using PTZ treatment at 5 mM. Both *kcnb1^+/-^* and *kcnb1^-/-^* larvae show a significant increase in distance travelled during the 30 min-recording after the chemical treatment as compared to WT zebrafish (N = 3 repeats; n = 27 ZF/genotype; one-way ANOVA with Bonferroni post-hoc test; ns: non-significant; +/- and -/- vs +/+ after PTZ treatment: **p<0,01; ****p<0,0001). **D.** RT-qPCR analysis of total *bdnf* at 6 dpf, a neurogenesis and epileptogenesis-related gene, before and after 5 mM PTZ treatment (N = 3; n = 30/sample; One-Way ANOVA with Bonferroni post-hoc test; ns: non significant; ****p<0,0001). Data are normalized to *ef1*α mRNA expression and *kcnb1^+/+^* non-treated fish are considered as the reference value (relative fold change = 1). The *kcnb1^-/-^* presents a tendency to an increased bdnf expression during the basal locomotor activity which is confirmed by a significant increased expression after chemically induced-seizures as compared to *kcnb1^+/+^ and Kcnb1^+/-^*. **E.** Whole-mount images of 6 dpf-zebrafish immunostained with anti-c-Fos, an acute neuronal activation marker, obtained 30 minutes after a basal activity period and a provoked-seizures period due to 5 mM PTZ treatment. The telencephalon was the major region activated after the chemical treatment for each genotype (n=8-11 ZF/condition, dorsal view, 3D reconstruction, scale bar: 50 µm, magnification 20x). **F.** Quantification of the number of c-Fos positive neurons normalized to the volume (µm^3^) of the telencephalon of 6 dpf larvae-zebrafish during their basal activity or chemically treated using the IMARIS V10.1.0 software (Oxford Instruments) (see **Supporting Information 3A**). The results did not show any difference in the number of activated neurons in both mutant conditions *(kcnb1^+/-^* and *kcnb1^-/-^*) as compared to the WT condition (n = 8-11 fish/condition; Mann Whitney test; ns: non-significant). **G.** Distribution (in %) of activated neurons according to the fluorescence intensity value of c-Fos in the telencephalon of fish at 6 dpf. Neuronal activation that was divided in four equal shares: low (0-25%), moderately low (25-50%), moderately high (50-75%) and high (75-100%) neuronal activation. The three non-treated conditions were presenting a similar distribution with a majority of low neuronal activation of c-Fos positive neurons. However, chemically treated *kcnb1^-/-^* zebrafish were presenting a global higher activation with a significant shift to a moderately low and moderately high neuronal activation (see **Supporting information 3B**). n = 8-11 fish/condition; Mann Whitney test; ns: non significant; *p<0,05; T: Telencephalon.

### 3.5. kcnb1 knock-out zebrafish model show spontaneous and provoked “epileptic”-like seizures associated with disrupted GABA regulation

Abnormal electrographic activity has been observed in zebrafish models with chemically provoked-epileptic seizures (26,31,34). We recorded local field potentials (LFPs) from the optic tectum of 6 dpf larvae during a 30-minute “basal activity” phase, followed by a 40 mM PTZ treatment, recorded for an additional 30 minutes **(Figure 5A-5B)**. Prior to PTZ treatment, *kcnb1^-/-^* larvae showed significantly increased spontaneous neuronal activity, reflected by the number of spikes, compared to WT **(Figure 5Ca-5D)**. In contrast, *kcnb1^+/-^* larvae did not exhibit spontaneous seizures, displaying a similar profile to untreated WT zebrafish **(Figure 5Ca-5D)**. In response to 40 mM PTZ exposure, WT larvae exhibited a significant increase in number of spikes compared to the non-treated condition **(Figure 5Ca-5D)**. The *kcnb1^+/-^*model showed a similar response profile, although the duration of seizure events was significantly longer than in PTZ-treated WT larvae **(Figure 5Ca-5E)**. Interestingly, *kcnb1^-/-^* larvae had a significantly higher number of PTZ-induced events, although event duration was comparable to WT-treated zebrafish **(Figure 5Ca-5E)**. This *kcnb1^-/-^* model also displayed various « epileptic »-like signals seen in *KCNB1*-related DEE patients, including polyspike discharges **(Figure 5Cb)**, ‘ictal’-like activity **(Figure 5Cc)**, and large amplitude spikes **(Figure 5Cd)**. To further confirm dysregulated neuronal activity in the *kcnb1* LOF model, we measured gamma-aminobutyric acid (GABA) and glutamate concentrations in the heads of 6 dpf zebrafish following the same protocol and exposed to 5 mM PTZ **(Figure 5A)**. During the pre-treatment period, GABA levels were similar in *kcnb1^+/-^*and WT larvae, while *kcnb1^-/-^* larvae showed a significant increase **(Figure 5F)**. Although PTZ treatment significantly increased GABA concentration in WT zebrafish, it did not alter GABA levels in the *kcnb1^+/-^* model, which remained low, or in the *kcnb1^-/-^*model, showing similar high concentration of GABA as WT-treated larvae **(Figure 5F)**. Glutamate levels were similar between all genotypes before PTZ exposure and remained unchanged in post-treatment **(Figure 5G)**. These results suggest that the *kcnb1^-/-^* model exhibits spontaneous ans chemically induced epileptiform-like electrographic activity, along with disrupted GABA regulation.

**Figure 5.**
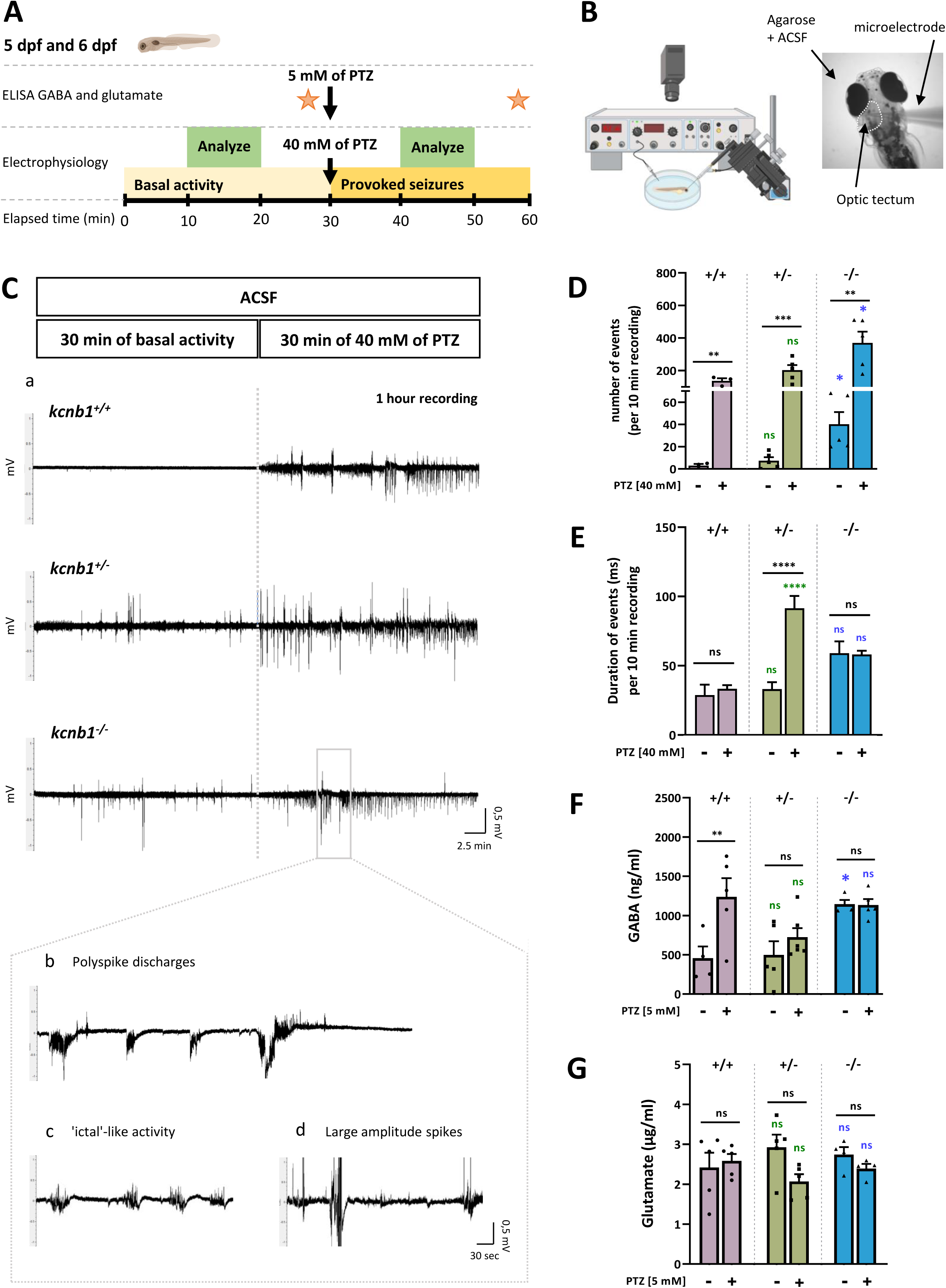
*kcnb1^-/-^*larvae show spontaneous and provoked epileptiform-like electrographic activity associated with disrupted GABA regulation. A and B. Schematic representation of the protocol applied for electroencephalographic recordings of 5 and 6 dpf zebrafish. Neuronal activity in the optic tectum of zebrafish was recorded by applying a protocol of 30 minutes of basal activity followed by 30 minutes of 40 mM PTZ treatment. 10 minutes in the middle of each period (basal activity and provoked seizures) was used to analyze the total number of negative spikes and duration of events for each genotype. Enzyme-linked immunosorbent assay (ELISA) was performed to quantify gamma-aminobutyric acid (GABA) and glutamate following a 30 minutes pre- and post-PTZ treatment at 5 mM. **C. (a)** Representative traces of electroencephalographic recordings in the optic tectum of WT and *kcnb1* knock-out larvae showing spontaneous and provoked epileptiform-like electrographic activity in the *kcnb1^-/-^* model characterized by **(b)** polyspike discharges, **(c)** ‘ictal’-like activity and **(d)** large amplitude spikes. **D-E.** Quantification of electrophysiological recordings by analyzing **(D)** the total number of negative spikes and **(E)** the duration of events, over 10 minutes as described in Figure 5A. *kcnb1^-/-^* larvae showed significantly increased spontaneous and provoked neuronal activity, reflected by a significant elevation of the number of spikes, although the duration of events was similar to the WT condition in pre- and post-PTZ treatment. *kcnb^+/-^* larvae presented similar profile as the WT condition in term of number of spikes but showed a signifiant increase of event duration (see **Table 3**; n = 3-5 ZF/genotype; unpaired t-test; ns: non significant; **p<0,01; ***p<0,001; ****p<0,0001). **F-G.** Quantification of **(F)** GABA and **(G)** glutamate by ELISA assays in the head of 6 dpf larvae following the 30-minutes pre- and post-5 mM PTZ treatment. *kcnb1^-/-^* zebrafish present significantly elevated GABA levels observed in both baseline and post-PTZ conditions conversely to *kcnb1^+/-^*larvae, compared to the WT line. We observed a lack of significant changes in glutamate levels in both *kcnb1* mutant models in pre- and post-PTZ conditions (N = 3-5 repeats; n = 50 heads/sample; unpaired t-test; ns: non significant; *p<0,05; **p<0,01). For figures 5D to 5G, colored statistic indications correspond to mutant conditions compared to the WT condition in pre- or post-PTZ treatment.

**Table 3:** Individual data of negative spikes number during electrophysiological recordings into the optic tectum of zebrafish. Neuronal activity of fish was recorded during 30 minutes of basal activity followed by 30 minutes of 40 mM PTZ treatment. A period of 10 minutes in the middle of each period (basal activity and provoked seizures) was used to analyze the number of negative spikes. The table shows a higher number of spikes for *kcnb1^-/-^* larvae in pre- and post-PTZ treatment as compared to WT fish. The following parameters were applied for each recording: Resistance: 2-7 MΩ; Low-pass Gaussian filter: 560 Hz; Data reduction: 10. n = 3-5 ZF/genotype.

| Condition | *kcnb1*<sup>+/+</sup> |  |  | *kcnb1*<sup>+/-</sup> |  |  |  |  | *kcnb1*<sup>-/-</sup> |  |  |  |  |
| --- | --- | --- | --- | --- | --- | --- | --- | --- | --- | --- | --- | --- | --- |
| Animal | 1 | 2 | 3 | 1 | 2 | 3 | 4 | 5 | 1 | 2 | 3 | 4 | 5 |
| Number of spikes in basal activity per 10 min recording | 0 | 5 | 4 | 12 | 6 | 17 | 2 | 0 | 68 | 20 | 21 | 66 | 26 |
| Number of spikes after 40 mM of PTZ per 10 min recording | 158 | 148 | 105 | 238 | 115 | 213 | 291 | 157 | 508 | 510 | 179 | 239 | 415 |

## 4. DISCUSSION

In this study, *kcnb1* knock-out in zebrafish results in early-onset phenotypes mimicking key features of *KCNB1*-related DEE. This LOF impacts neuronal functions in particular inhibitory pathways in developemental brains.

### 4.1. kcnb1 expression in the CNS

The study of kcnb1 expression revealed the presence of the protein across various regions of the zebrafish CNS, including the diencephalon, midbrain, telencephalon, and hindbrain, consistent with previously reported *in situ* hybridization of *kcnb1* by Shen and collaborators (2016). This broad expression supports previous findings that kcnb1 is crucial for maintaining neuronal excitability, starting from 19 hpf (35). Notably, the presence of kcnb1 in multiple cell subtypes suggests its diverse functional roles, ranging from neuronal signaling to potential involvement in glial cell function and neuroinflammation in zebrafish.

### 4.2. kcnb1 knock-out and developmental consequences

Despite a significant reduction in *kcnb1* transcript levels in both heterozygous and homozygous mutants, the protein expression was similar between mutant models and the WT condition, suggesting a potential genetic compensation. We observed no major morphological, brain anatomical abnormalities or impaired survival rates in *kcnb1^-/-^* zebrafish during the early development. This finding is consistent with clinical observations in *KCNB1*-related DEE patients, where normal brain morphology is reported in most cases despite severe neurological symptoms (9). This suggests that *kcnb1* LOF does not impact brain structural development, although it may influence more neuronal function aspects.

### 4.3. Altered behavioral phenotypes and sensorimotor dysregulation

Behavioral assays revealed that *kcnb1^-/-^* zebrafish exhibit significant locomotor impairments, including hyperactivity, altered swimming patterns, and exaggerated responses to sensory stimuli such as light and sound. Indeed, we showed that *kcnb1*^+/-^ and *kcnb1*^-/-^ embryos had reduced tail coiling activity from 25 hpf to 31hpf. This early stereotyped behavior is supported by synchronized spinal locomotor circuits (39). Previous reports on models of epilepsy have revealed a corkscrew-like trajectory swimming characteristic of epileptogenic-like activity (22). We found a similar circular pattern of swim trajectories in *kcnb1* mutant models at 48 hpf, but revealed a huge variability within the same genotype. These results might reflect the variable spectrum of behavioral and cognitive impairments observed in patients (12). Furthermore, *kcnb1*^-/-^ mutants exhibited increased response of locomotor activity to a light-dark stimulus as compared to *kcnb1*^+/-^ and WT lines, but presented a decreased locomotor activity after a succession of audio stimuli. This last result is in correlation with the data obtained by Jedrychowska and collaborators (2021). These phenotypes closely mimic the motor dysfunctions observed in patients with *KCNB1* mutations, such as ataxia and hyperactivity and suggest a dysregulation of sensorimotor pathways.

### 4.4. Seizure susceptibility and electrophysiological abnormalities

The seizure-like behavior in *kcnb1*^+/-^ and *kcnb1*^-/-^ models is supported by their increased locomotor activity characterized by fast circles trajectories induced by 5 mM PTZ compared to wild-type. These results are concomitant with the stage II of the “seizure-like behavior score” described by Baraban and collaborators (2005). Our electrophysiological recordings further demonstrated that *kcnb1^-/-^* zebrafish exhibit spontaneous and chemically induced epileptiform seizure-like activity. The absence of spontaneous seizures in the *kcnb1^+/-^* line could reflect the existence of functional compensatory pathways masking the phenotypic features due to reduced levels of *kcnb1* mRNA but similar protein expression as WT larvae. Epileptiform seizure-like activity was confirmed by c-Fos acute neuronal hyperactivation in the telencephalon of *kcnb1*^-/-^ zebrafish. Moreover, the quantification of brain-derived neurotrophic factor *(bdnf)* mRNA in larvae, a neurotrophin crucial for brain development and synaptic plasticity, revealed a significant upregulation of its expression level within the *kcnb1*^-/-^ line. This finding indicates deficits in neurogenesis within the developing brain of *kcnb1*^-/-^ zebrafish. Similar results were obtained by identifying *c-Fos* and *bdnf* as genes associated with seizures in zebrafish, with increased expression levels observed after exposure to 20 mM PTZ 20mM (40). These findings are particularly relevant to understanding the pathophysiology of DEEs, where patients often present with refractory seizures and abnormal electroencephalogram patterns.

### 4.5. Neurotransmitter dysregulation in kcnb1^-/-^ zebrafish

Our study also highlights the dysregulation of GABA in *kcnb1^-/-^*zebrafish, with significantly elevated GABA levels observed in both baseline and post-PTZ conditions. This disrupted GABAergic signaling likely contributes to the observed seizure phenotypes, as GABA is a key inhibitory neurotransmitter involved in maintaining the balance of excitatory and inhibitory signals in the CNS. The lack of significant changes in glutamate levels suggests that *kcnb1* LOF primarily affects inhibitory pathways, leading to an imbalance that favors neuronal hyperexcitability.

## 5. CONCLUSION

Our investigation suggests that the *kcnb1* loss-of-function zebrafish model provides a valuable model to reproduce key features of *KCNB1*-related DEE, including early behavioral disturbances, increased-susceptibility to epileptic seizures and neurotransmitter dysregulation. Notably, this model could be used for discovering of new therapeutic compounds that may improve the long-term prognosis of individuals with *KCNB1*-related DEE.

## AUTHOR CONTRIBUTIONS

Lauralee Robichon designed and conceptualized the study, performed the experiments, analyzed and interpreted the data, and wrote the manuscript. Claire Bar designed some parts of the study, performed some behavioral assays and wrote the manuscript. Anca Marian performed some behavioral assays and was responsible for technical support. Lisa Lehmann performed some locomotor activity experiments and analyzed some data. Solène Renault was responsible for technical support. Edor Kabashi supervised the work, revised the paper and approved the manuscript. Sorana Ciura supervised the work, revised the paper and approved the manuscript. Rima Nabbout supervised the work, revised the paper and approved the manuscript. All authors revised and approved the final version of the manuscript.

## ACKNOWLEGMENTS

This work was supported by grants from the Agence Nationale de la Recherche under “Investissements d’avenir” program (ANRl10IAHUl01), the Fondation Bettencourt Schueller (Rima Nabbout and Claire Bar), the Ligue Française Contre l’Épilepsie (Claire Bar), the ERC Consolidator Grant (Edor Kabashi). Rima Nabbout and Lauralee Robichon are supported by the Chair Geen-DS funded by FAMA fund hosted by Swiss Philanthropy Foundation. Lauralee Robichon is recipient of a grant from the Fondation pour la Recherche Médicale (grant number: PLP202009012460, 2021). The work was supported by the “Association KCNB1 France”.

We are grateful of the team of Dr. Vladimir Korzh (International Institute of Molecular and Cell Biology of Warsaw, Poland) for kindling provided us the *kcnb1* knock-out transgenic zebrafish line. We appreciate the help of the LEAT Zebrafish facilities and Cell Imaging Facility of Imagine Institute for respectively fish maintenance and expert technical help. The help of Nicolas Goudin, responsible of the Image analysis center of SFR Necker (Paris, France) was highly appreciated for immunofluorescence quantification. We thank DSHB for the antibodies used for immunohistochemistry: 3A10 was deposited to the DSHB by Jessell, T.M. / Dodd, J. / Brenner-Morton, S. (DSHB Hybridoma Product 3A10), PCRP-OLIG2-1E9 was deposited to the DSHB by Common Fund – Protein Capture Reagents Program (DSHB Hybridoma Product PCRP-OLIG2-1E9).

## DISCLOSURE

None of the authors has any conflict of interest to disclose.

We confirm that we have read the Journal’s position on issues involved in ethical publication and affirm that this report is consistent with those guidelines.

## SUPPORTING INFORMATIONS

**Supporting information 1. Brain anatomy and neuronal circuits are not disrupted in the *kcnb1* loss-of-function zebrafish model. A.** Horizontal section of WT zebrafish expressing cells (DAPI, blue) labelled with anti-kcnb1 (green), showing a large expression of the protein in various regions of the central nervous system (CNS) at 6-day post-fertilization (dpf) (scale bar 50 µm; magnification10x; n = 3-4 fish/section). **B.** Horizontal sections of WT zebrafish expressing cells (DAPI, blue), kcnb1 (green) and Ankyrin G expressed at the axonal initial segment of neurons (AnkG, purple, scale bar 30 µm; magnification 20x; n = 3-4 fish/section). The image demonstrate the colocalization between kcnb1 and AnkG, represented in white and marked by arrows, indicating the presence of kcnb1 in neurons. Colocalization was determined using Z-stack projection on IMARIS V10.1.0 software (Oxford Instruments). In figures, CNS regions are delimited by dotted lines. **C.** Transversal slices of 48 hpf and 4 dpf larvae expressing cells (DAPI, blue) labelled with anti-HuC (red), a pan-neuronal marker, showing no major difference of anatomical brain regions between mutants and wild-type fish (scale bar: 50 µm; magnification 20x; n = 3-4 fish/section). **D.** Whole-mount images of embryonic (48 hpf) and larvae (6 dpf) zebrafish immunostained with anti-3A10 marker to identify Mauthner cells (dorsal view; 3D reconstruction; scale bar: 50 µm; magnification 20x; n = 3-5 fish/section). **E.** Quantification of Mauthner cell body length (M-cell) of fish at 48 hpf between the two extremities as represented with the arrow. The average value of the two M-cell bodies of each fish was used for quantification. Both *kcnb1* knock-out zebrafish models (*kcnb1^+/-^* and *kcnb1^-/-^*) do not present difference in the length of M-cell body at 48 hpf as compared to the WT condition (n = 3 - 5 ZF/genotype; one-Way ANOVA with Bonferroni post-hoc test; ns: non-significant). C: Cerebellum; D: Diencephalon; DAPI: 4,6ldiamidinol2lphenylindole; E: Eyes; H: Hindbrain; kcnb1: Potassium voltage-gated channel subfamily B member 1; M: Midbrain; OT: Optic tectum; SC: Spinal cord; T: Telencephalon.

**Supporting information 2: *kcnb1* mutant zebrafish models present a premotor impairment and variability in the swimming behavior. A.** Locomotor activity heatmap reflecting the average number of movements per minute of embryos in their chorion between 24 hpf and 36 hpf. **B.** A significant decrease in general spontaneous movements (coiling and twitching) of *kcnb1* mutant models (*kcnb1^+/-^* and *kcnb1^-/-^*) was observed as compared to the WT condition (*kcnb1^+/+^*) during the whole first day of development (N = 3 repeats; n = 80-104 ZF/genotype; one-way ANOVA with Bonferroni post-hoc test; ns: non-significant; *p<0,05; **p<0,01; ****p<0,0001). **C.** Within the same genotype, the swimming behavior of fish has been divided into two distinct phenotypes: severe (at least two swim circles, severe phenotype, SP) and mild (other trajectories, mild phenotype, MP) during the 2^nd^ dpf. The subdivision of both swimming behavior (severe and mild phenotype) per genotype is represented as traces of individual swimming episodes showing a huge variability of trajectories (4 trajectories/phenotype). 30,68% of *Kcnb1^+/-^* fish and 52% of *kcnb1^-/-^* fish were identified with a severe phenotype (see **Table 2**). **D-F.** The variability between both swimming phenotypes was quantified following different parameters including **(D)** the distance swam, **(E)** the velocity and **(F)** the time spent in motion. The results show similar values between both mutant conditions presenting a mild phenotype as compared to the WT condition. Nevertheless, fish with a severe phenotype present a significant increase of the studied parameters for both genotypes (*kcnb1^+/-^* and *kcnb1^-/-^*). N = 3 repeats; n = 50-88 ZF/genotype; one-way ANOVA with Bonferroni post-hoc test; **p<0,01; ***p<0,001; ****p<0,0001.

**Supporting information 3: Shift of c-Fos positive-neuron intensity distribution from a global low neuronal activation to a higher neuronal activation in the telencephalon region of *kcnb1^-/-^* fish at 6 dpf after chemically provoked-seizures. A.** Whole larvae-fish at 6 dpf were fixed 30 minutes after a basal activity period and a provoked-seizures period due to PTZ treatment at 5 mM followed by the labelling of c-Fos, an acute neuronal activation marker. The number of c-Fos positive-neurons and the fluorescence intensity value of c-Fos per neuron were quantified using the software IMARIS by analyzing the telencephalon, the major region activated after the chemical treatment. The contour of the region of interest has been designed manually to obtain the volume (µm^3^) of the telencephalon, neurons were then identified automatically within the region following identical parameters for spot detection between conditions and visualized in 3 dimensions to avoid false negative spots. Values were normalized according to the volume of the telencephalon (see Figure 4E**-4F**; n = 8-11 fish/condition). **B.** Distribution (in %) of c-Fos positive-neurons according to the fluorescence intensity value of c-Fos in the telencephalon of fish at 6 dpf, reflecting the level of neuronal activation that was divided in four equal shares (from 0 to 100%): low (0-25%), moderately low (25-50%), moderately high (50-75%) and high (75-100%) neuronal activation. During the basal activity, the distribution of activated neurons is similar between each conditions with a majority of neurons presenting a low neuronal activation. However, a shift to a moderately low and moderately high neuronal activation was observed for each condition after a PTZ treatment, with a global higher activation in the *kcnb1^-/-^* condition (see Figure 4G). n = 8-11 fish/condition; T: Telencephalon.

