## Supplementary figures and images for "*kcnb1* loss-of-function in zebrafish causes neurodevelopmental and epileptic disorders associated with GABA dysregulation"

### supplementary data 1

#### Supporting information 1.

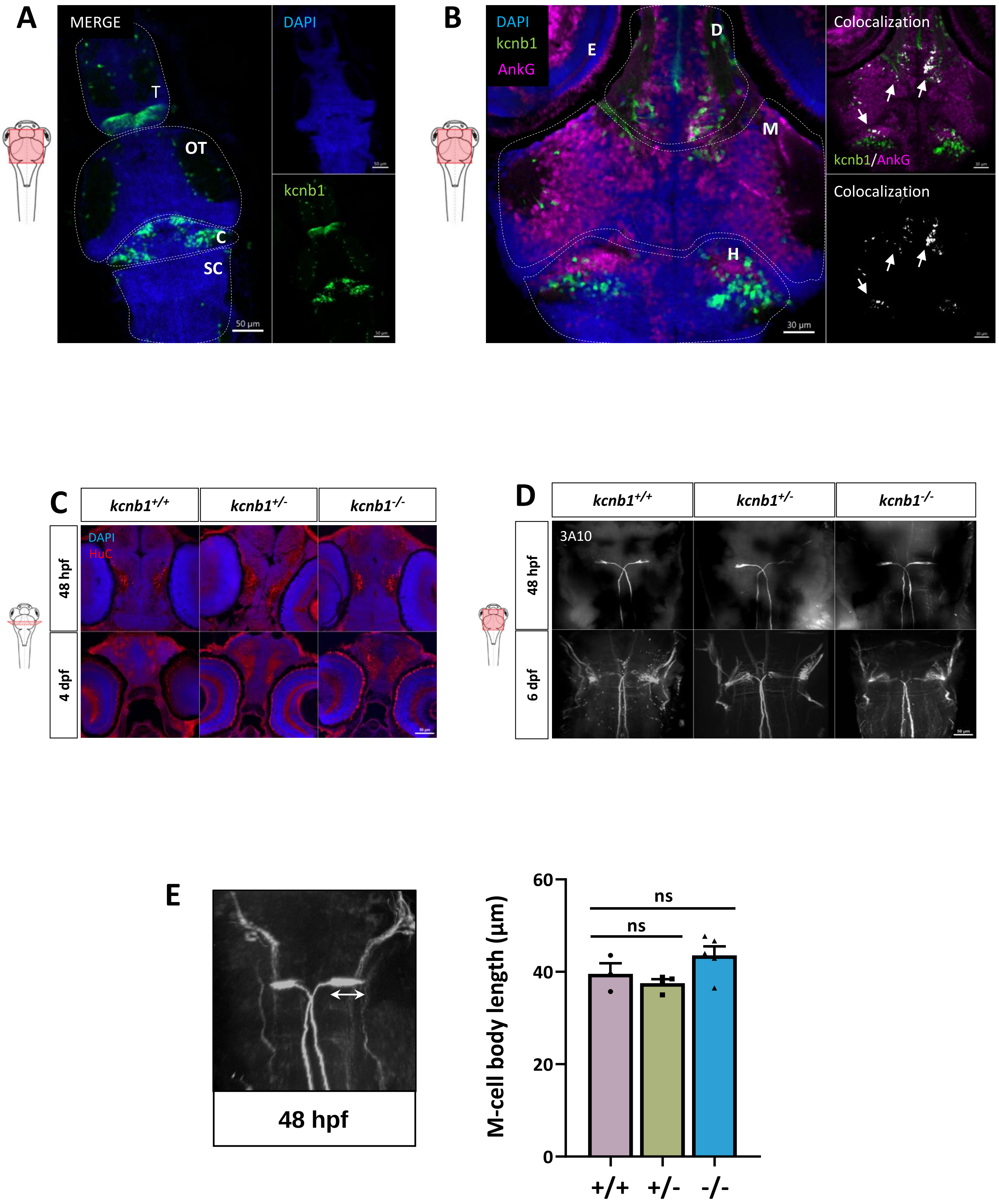

### supplementary data 2

#### Supporting information 2.

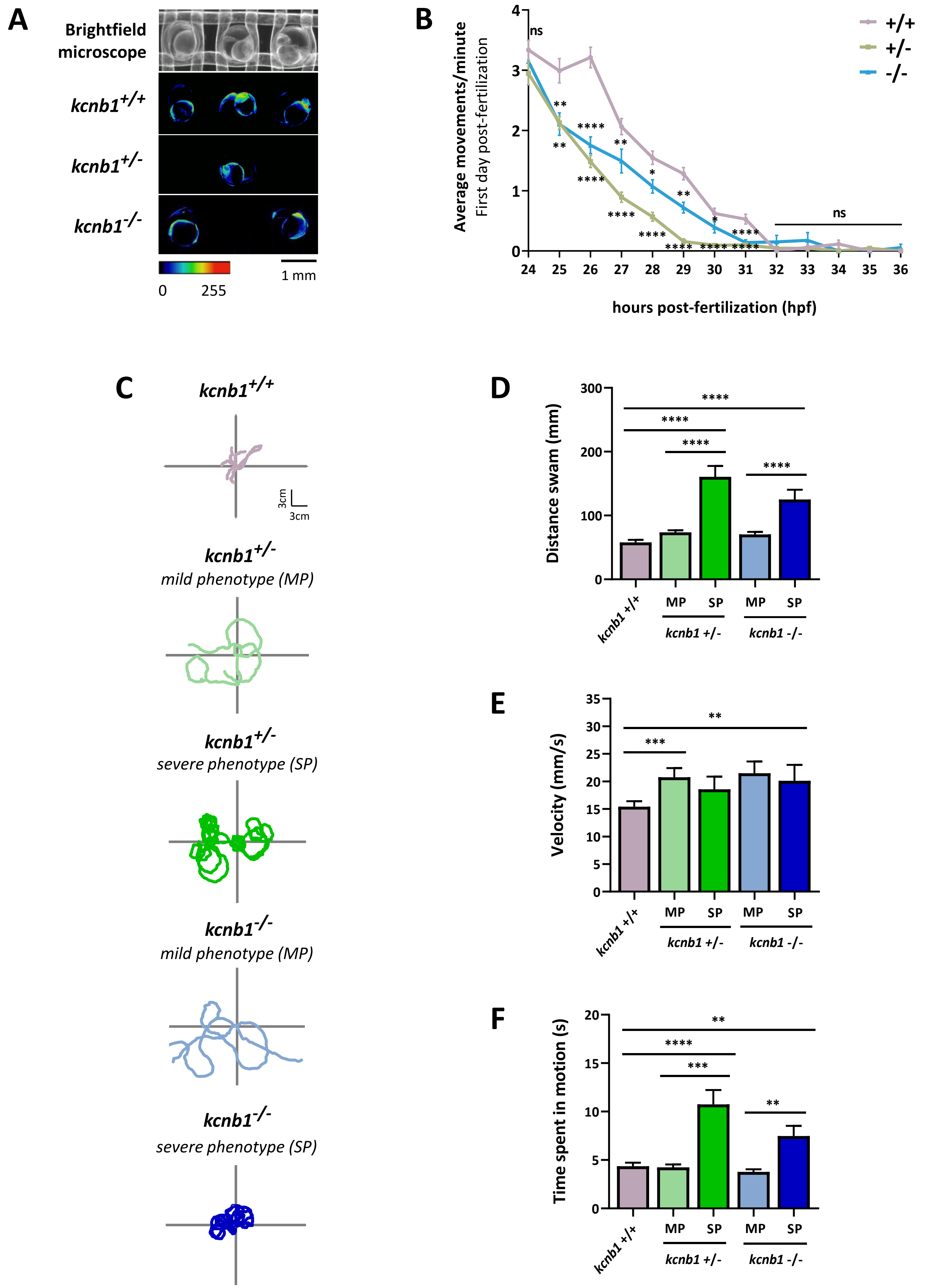
