## supplementary data 3 for "*kcnb1* loss-of-function in zebrafish causes neurodevelopmental and epileptic disorders associated with GABA dysregulation"

### Supporting information 3.

**A**

1. Region delimitation
2. Area detection
3. Dots detection inside the area
4. 3D dots identification

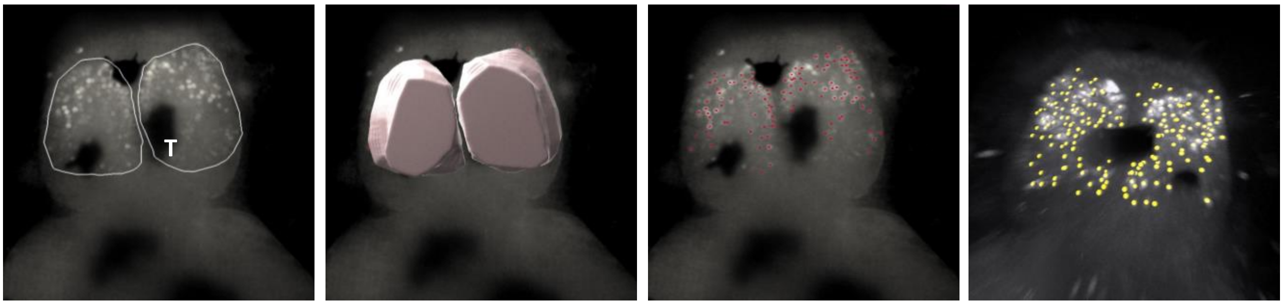


**B** **Distribution (in %) of activated neurons according to the c-Fos fluorescence intensity in the telencephalon region of 6 dpf-zebrafish**

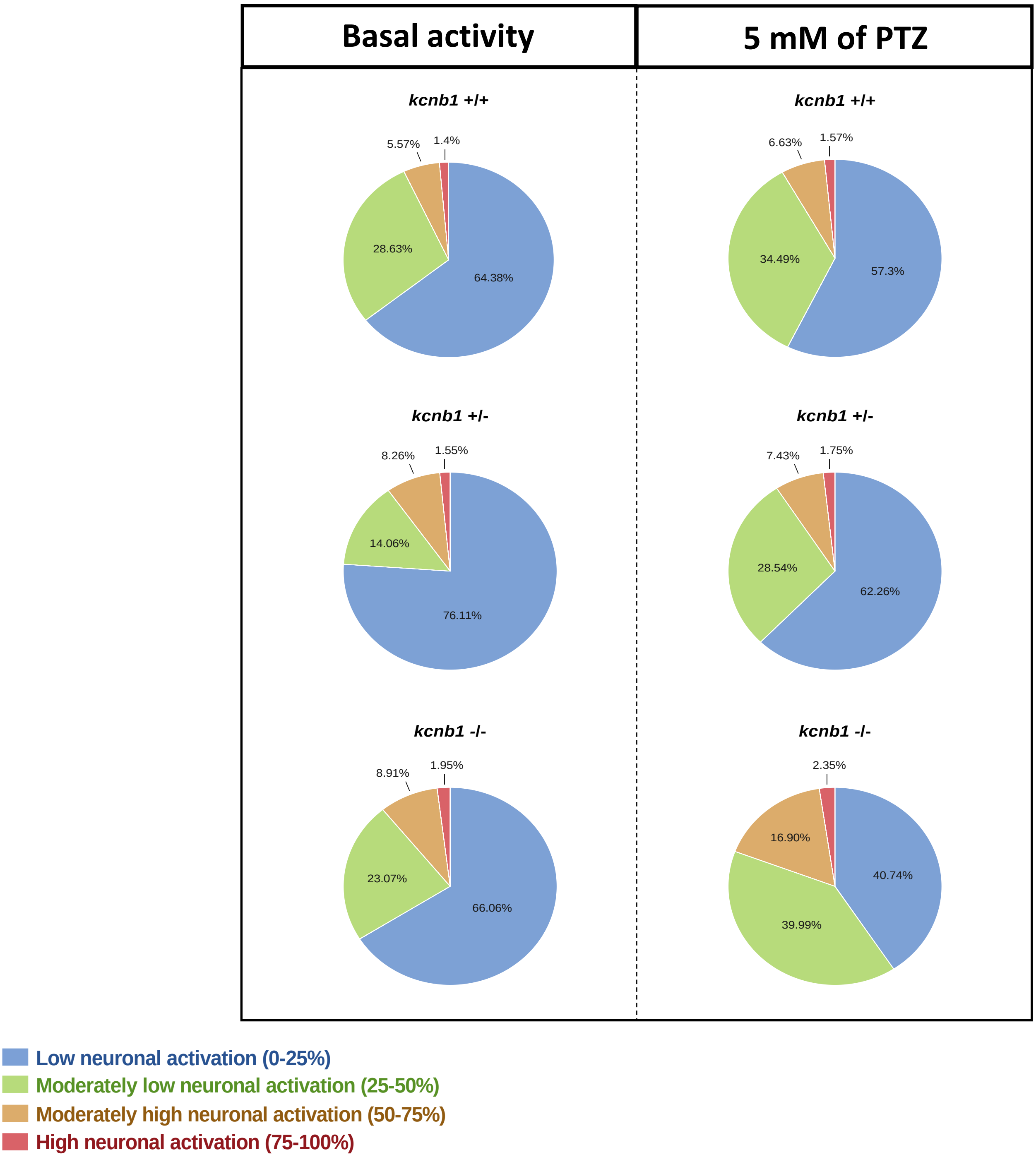
